# Structural requirements of the phytoplasma effector protein SAP54 for causing homeotic transformation of floral organs

**DOI:** 10.1101/772699

**Authors:** Marc-Benjamin Aurin, Michael Haupt, Matthias Görlach, Florian Rümpler, Günter Theißen

## Abstract

Phytoplasmas are intracellular bacterial plant pathogens that cause devastating diseases in crops and ornamental plants by the secretion of effector proteins. One of these effector proteins, termed SECRETED ASTER YELLOWS-WITCHES’ BROOM PROTEIN 54 (SAP54), leads to the degradation of a specific subset of floral homeotic proteins of the MIKC-type MADS-domain family via the ubiquitin-proteasome pathway. In consequence, the developing flowers show the homeotic transformation of floral organs into vegetative leaf-like structures. The molecular mechanism of SAP54 action involves physical binding to the keratin-like K-domain of MIKC-type proteins, and to some RAD23 proteins, which translocate ubiquitylated substrates to the proteasome. The structural requirements and specificity of SAP54 function are poorly understood, however. Here we report, based on biophysical and molecular biological analyses, that SAP54 folds into α-helical structures. We also show that the insertion of helix-breaking mutations disrupts correct folding of SAP54, which interferes with the ability of SAP54 to bind to its target proteins and to cause disease phenotypes *in vivo*. Surprisingly, dynamic light scattering data together with electrophoretic mobility shift assays suggest that SAP54 preferentially binds to multimeric complexes of MIKC-type proteins rather than to dimers or monomers of these proteins. Together with literature data this finding suggests that MIKC-type proteins and SAP54 constitute multimeric α-helical coiled-coils, possibly also involving other partners such as RAD23 proteins. Our investigations clarify the structure-function relationship of an important phytoplasma effector protein and thus may ultimately help to develop treatments against some devastating plant diseases.

**SIGNIFICANCE STATEMENT:** Phytoplasmas are bacterial plant pathogens that cause devastating diseases in crops and ornamental plants by the secretion of effector proteins such as SAP54, which leads to the degradation of some floral homeotic proteins. Our study clarifies the structural requirements of SAP54 function and illuminates the molecular mode of interaction and thus may ultimately help to develop treatments against some devastating plant diseases.

## Introduction

Phytoplasmas (genus ‘*Candidatu*s Phytoplasma’) are mycoplasma-like bacterial symbionts of plants belonging to the class Mollicutes. Among pathogens they have a unique life cycle, as they invade and replicate in organisms of two different kingdoms, namely plants and animals (specifically, insects belonging to plant hoppers, leaf hoppers and Psyllids) (Sugio et al., 2011b). Inside plants, phytoplasmas live almost exclusively within the phloem, whereas in insects they colonize most major organs including the salivary glands from which the bacteria are transmitted to new plant hosts during feeding (Hogenhout et al., 2008; Sugio et al., 2011b). Phytoplasmas are highly derived organisms, in that they have lost their cell wall and went through massive genome reduction (reviewed by Christensen et al., 2005; Firrao et al., 2007; Hogenhout et al., 2008). A phytoplasma infection may induce severe changes in plant morphology via modification of developmental processes. Morphological changes include the induction of witches’ brooms (tight clustering of branches), phyllody (full or partial homeotic transformation of floral organs into leaf-like structures), virescence (green coloration of floral organs), purple top (reddening of leaves and stems), phloem necrosis, stunting of growth habit, and general yellowing of plants (Lee et al., 2000; Zhang et al., 2004). Since the plant hosts of phytoplasmas include many economically important crops, including fruit trees, and ornamental plants, these symptoms lead to economically devastating damages each year worldwide (reviewed by Bertaccini, 2007; Lee et al., 2000; Liu et al., 2017). Since infected plants allow the replication of phytoplasmas but are frequently not able to reproduce themselves anymore they were termed ‘zombie plants’ (Du Toit, 2014).

There is evidence that phytoplasmas induce most of the developmental and morphological changes by secreting virulence factors, termed effector proteins, into the phloem of the host plant via the bacterial Sec secretion system (Bai et al., 2009; Kakizawa et al., 2001). Effectors are small peptides characterized by the presence of a signal peptide required for secretion that have been shown to reprogram plant development in several ways (Bendtsen et al., 2004; Hoshi et al., 2009; Sugio et al., 2011a). Owing to their small size, the effectors are able to travel systemically through the plant and reach tissues distant to the phloem (Bai et al., 2009; Hoshi et al., 2009; Sugio et al., 2011b).

An particularly intriguing example is the interaction of the effector protein SECRETED ASTER YELLOWS-WITCHES’ BROOM PROTEIN 54 (SAP54) from *Candidatus* Phytoplasma asteris strain Aster Yellows Witches’ Broom (AY-WB), with MIKC-type MADS-domain transcription factors (MIKC-type MTFs) of the host plants (MacLean et al., 2014; MacLean et al., 2011; Maejima et al., 2014). MTFs control a plethora of plant developmental processes, in angiosperms ranging from root to flower and fruit development (reviewed by Smaczniak et al., 2012a; Theißen et al., 2016). Among the 45 different MIKC-type MTFs being present in the model plant *Arabidopsis thaliana,* only a subset is specifically bound by SAP54, most of which are involved in the control of floral meristem and organ identity (MacLean et al., 2014). SAP54 mediates the degradation of the targeted proteins via the ubiquitin-proteasome pathway (UPP) of the host plant (MacLean et al., 2014; Maejima et al., 2014; Maejima et al., 2015). According to one model, SAP54 binds the MIKC-type MTFs in a complex together with some RADIATION SENSITIVE 23 (RAD23) proteins, which play a central role in targeting ubiquitylated proteins for proteasomal degradation (MacLean et al., 2014; Maejima et al., 2014; Maejima et al., 2015). Since the degraded MTFs are required for the specification of organ identity during flower development, the resulting floral structures show the development of vegetative leaf-like structures rather than floral organs proper.

Within MIKC-type MTFs, SAP54 has been shown to specifically bind to the so called keratin-like domain (K-domain) (MacLean et al., 2014). The K-domain is shared by all MIKC-type MTFs and mediates protein-protein interactions that allow these proteins to form different homo- and heteromeric dimers and tetramers (Kaufmann et al., 2005; Melzer and Theißen, 2009; Melzer et al., 2009; Puranik et al., 2014; Smaczniak et al., 2012b). The amino acid sequence within the K-domain follows a characteristic pattern of regularly spaced hydrophobic and charged residues that repeats every seven amino acids (Ma et al., 1991; Riechmann et al., 1996). Such heptad repeat patterns, of the form [abcdefg]_n_ with ‘a’ and ‘d’ positions being mainly occupied by hydrophobic amino acids, are typical for protein α-helices that are able to form coiled-coil structures, a common and well-studied class of protein-protein interaction domains (reviewed by Lupas and Gruber, 2005; Mason and Arndt, 2004). According to X-ray crystallographic data of the MIKC-type MTF SEPALLATA3 (SEP3) from *A. thaliana*, which is a target of SAP54, the K-domain folds into two coiled-coils separated by a short kink (Puranik et al., 2014). As the amino acid sequence within the K-domain is highly conserved among MIKC-type MTFs it appears very likely that the K-domains of most MIKC-type MTFs fold in a structure very similar to that determined for SEP3 (Rümpler et al., 2018).

Based on *in silico* structure prediction we previously hypothesized that SAP54 folds into a structure that is very similar to that of the K-domain of MIKC-type MTFs; we also suggested that the mode of interaction between SAP54 and targeted MIKC-type MTFs resembles and mimics the one mediating protein-protein interactions among MIKC-type MTFs itself (Rümpler et al., 2015). Here, we experimentally tested these hypotheses *in vitro*, *in vivo* (yeast) and *in planta* using biophysical and mutational analyses, as well as protein-protein interaction studies. Our present data here strongly corroborate the view that SAP54 indeed folds into two α-helices that are capable of forming coiled-coils and that are separated by a short interhelical region - a structure remarkably similar to that of the K-domain of MIKC-type MTFs. Dynamic light scattering data and electrophoretic mobility shift assays (EMSA) suggest that this structure allows SAP54 to target MIKC-type MTFs via forming multimeric α-helical coiled-coils.

## RESULTS

### *In silico* predictions suggest that SAP54 folds into α-helices capable of forming coiled-coils

To get a first impression of the structural properties of SAP54 of phytoplasma AY-WB and its homologs we compiled a sequence collection of 28 SAP54-like proteins from a diverse set of phytoplasma strains based on BLAST searches in the non-redundant NCBI nucleotide and protein collections, and in whole genome data. The vast majority of sequences aligned without any gaps (i.e. without potential insertions or deletions). With an average sequence identity of ∼76 % the sequences appeared to be overall very similar (Figure 1a). Merely sequences from phytoplasma strains belonging to the 16SrII and the 16SrXII group deviate from the majority of sequences, especially within the first 15 amino acids of the secreted part and within the centre of the protein. In addition, six of the sequences appear truncated at their C-termini. Inspection of the coding sequences of the respective genes revealed that either a nonsense mutation (strains 284/09 and NJAY), single or double nucleotide insertions resulting in a frame shift and a premature stop codon (strains NTU2011, PYR, and WBD), or an incomplete genomic DNA sequence (strain MA1) caused the truncated sequences. Based on the aligned SAP54-like sequences we performed secondary structure predictions applying different programs. All secondary structure prediction programs consistently predicted a high α-helical content spanning almost the entire protein (Figure 1b). Only the first 20 amino acids at the N-terminus are presumably unstructured. The (at least) two putative helices were predicted to be separated by a turn or bend located in the centre of the protein. According to PCOILS the predicted helices show a continuous heptad repeat pattern and consequently a high probability to form coiled-coils. We further investigated the structure of SAP54 by conducting different 3D structure predictions. QUARK Online, PSIPRED, and SWISS-MODEL produced very similar 3D structures according to which SAP54 folds into at least two coiled-coils that are separated by an interhelical region (Figure 1c).

**Figure 1.**
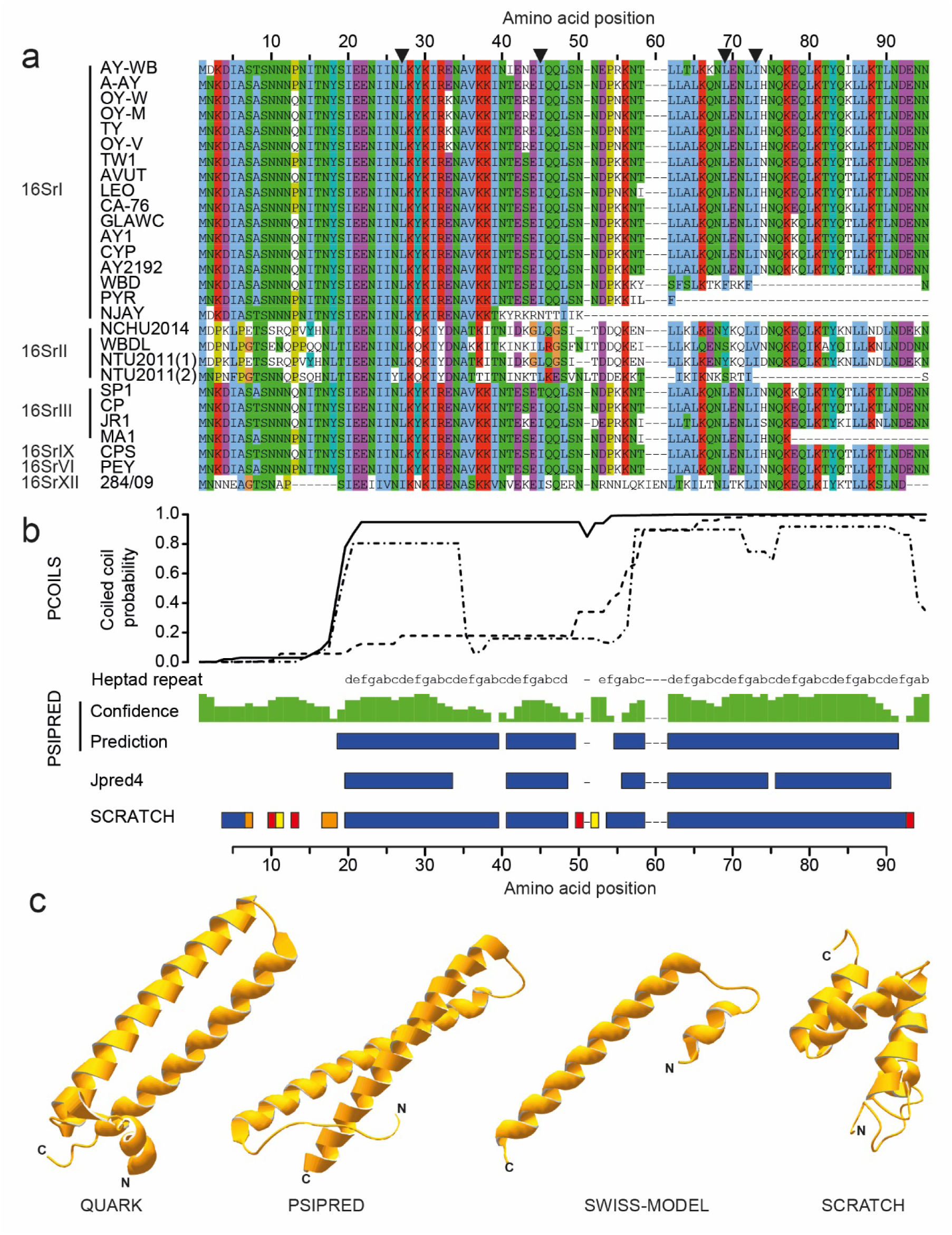
Sequences and predicted structural features of SAP54-like proteins (phyllogens). (a) Amino acid alignment of the secreted part of SAP54-like sequences from different phytoplasma strains. Sequences were aligned with MAFFT applying L-INS-I mode. Black triangles on top of the alignment highlight the amino acid positions of SAP54 from AY-WB that were substituted by site directed mutagenesis. SAP54 homologues PHYL1_OY_ from Onion Yellows phytoplasma and PHYL1_PnWB_ from Peanut witches’ broom phytoplasma, that have been recently crystallized can be found under the synonyms “OY-W” and “NTU2011 (1)”, respectively. (b) Secondary structure predictions based on the amino acid sequence of SAP54 from AY-WB (using Jpred4 and SCRATCH Protein Predictor) and the multiple sequence alignment shown in (a) (using PCOILS, PSIPRED). PCOILS: Dotted, dashed, and solid lines show the coiled-coil probability of the amino acid alignment in (a) for a sliding window size of 14, 21, and 28 amino acids, respectively. PSIPRED, Jpred4, and SCRATCH: Colored boxes indicate high probability for the formation of an α-helix (blue), extended strand (orange), turn (red), or bend (yellow), respectively. The green bar plot depicts the confidence of the PSIPRED prediction. (c) 3D structure predictions for SAP54 from AY-WB using QUARK Online, PSIPRED, SWISS-MODEL and SCRATCH Protein Predictor. The SWISS-MODEL prediction only comprises the C-terminal part of SAP54 (residues 43-91).

### SAP54 shows high α-helical content

In order to experimentally address the presence of the predicted α-helical elements of SAP54 we heterologously expressed SAP54 in *Escherichia coli*, purified it and recorded circular dichroism (CD) spectra. The CD spectrum recorded for SAP54 exhibited two minima at 222 and 208 nm, respectively, as well a distinct maximum near 195 nm. Both features constitute hallmark spectral properties of α-helical proteins (Figure 2a) (Kelly et al., 2005; Ranjbar and Gill, 2009). Estimated from the residual ellipticity at 222 nm, SAP54 showed an α-helical content of ∼71 % at the applied conditions (Table 1). In contrast, no obvious β-strand content and only minor irregular structure features were determined, likely to indicate presence of loops or disordered regions. In order to monitor the temperature dependence of the folding state of SAP54, its molar ellipticity at 200 nm and 222 nm was determined beginning with 20°C in 5°C intervals up to 90°C and depicted according to Uverksy (Uversky, 2002). This allows assessment of the folding state of a protein for the chosen temperatures. At 20°C, SAP54 appears in the region for folded proteins (Figure S1) with gradual loss of secondary structure content as the temperature is increased. At 80°C to 90°C, SAP54 appears to adopt a ‘molten globule’ state, which would suggest, that it does not entirely denature even at 90°C (Figure S1, Table S1). Reducing subsequently the temperature back to 20°C, SAP54 renatures to a certain degree, albeit not to its initial folded state (Figure S1).

**Figure 2.**
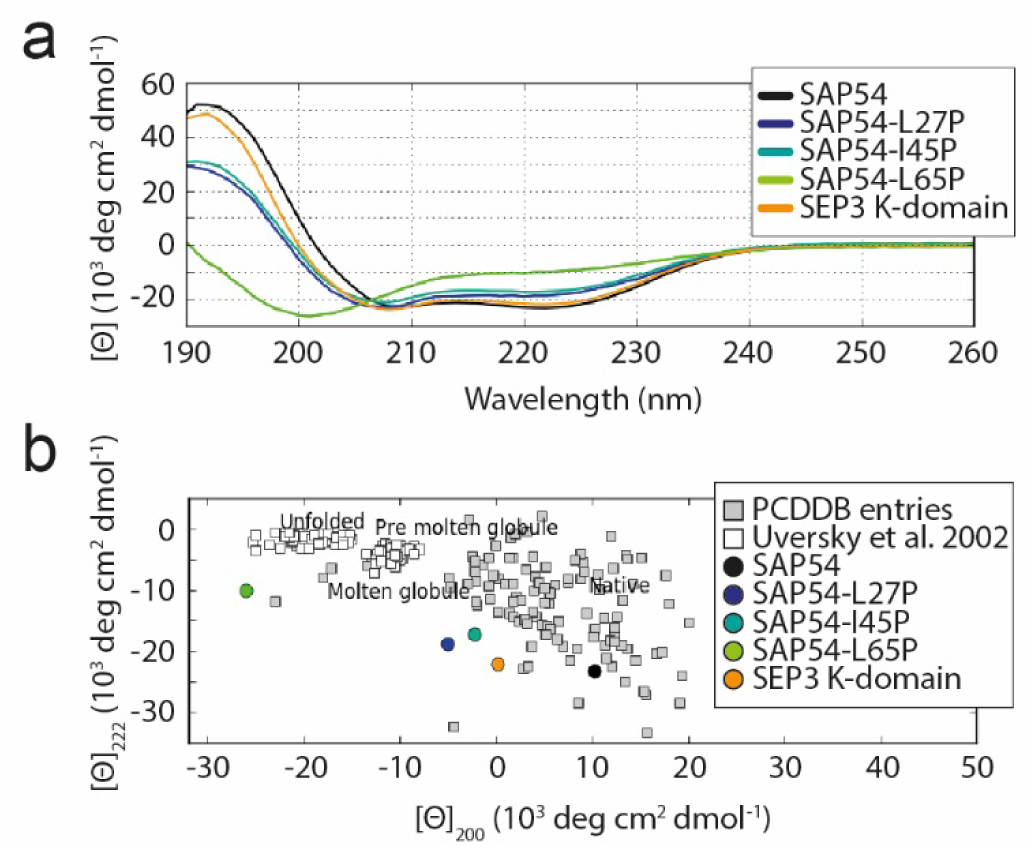
CD spectra of SAP54-wt, its mutated versions and the K-domain of SEP3. (a) CD spectra recorded on JASCO J-715 spectropolarimeter for SAP54-wt, SAP54-L27P, SAP54-I45P and SAP54-L65P and the K-domain of SEP3 at 20°C in 20 mM sodium phosphate buffer, pH 8.0, wavelength range of 190-280 nm at 1 nm intervals and a speed of 20 nm/min using a cuvette of 1 mm path length. Protein concentrations were: SAP54-wt: 20.7 µM; SAP54-L27P: 6.5 µM, SAP54-I45P: 10.6 µM; SAP54-L65P: 9.9 µM. Each spectrum represents an accumulation of five background corrected spectra. (b) Placement of the examined proteins in the Uversky plot, which allows a categorization of the folding state of the different proteins. Graphics generated with the help of CAPITO.

**Table 1.** Secondary structure content of SAP54-wt, single amino acid substitution mutants and the K-domain of SEP3. Values were calculated with CAPITO based on the recorded CD spectra in sodium phosphate buffer at pH 8.0 at room temperature. SAP54-wt values represent means based on 3 biological replicates.

| Proteins | % α-helix | % β-strand | % irregular |
| --- | --- | --- | --- |
| SAP54 | 71 ± 9 | 1 ± 0 | 29 ± 2 |
| SAP54-L27P | 42 | 1 | 55 |
| SAP54-I45P | 53 | 1 | 52 |
| SAP54-L65P | 7 | 1 | 82 |
| SEP3 K-domain | 65 | 1 | 46 |

### Truncation of putative helices results in a loss of SAP54 interaction ability

Our *in silico* analyses suggest that the high α-helical content of SAP54 probably involves two helices that are separated by a turn or bend located near the center of the protein (Figures 1b and c). Thus we investigated whether deletions of either of the putative helices interferes with the interaction capabilities of SAP54 with its known MIKC-type MTF targets. We generated truncated versions of SAP54 in which the first predicted helix (amino acids 1-48 of the mature SAP54, Figure 3a), the putative turn region (amino acids 49-58), or the second predicted helix (amino acids 59-91) were missing, and tested the interaction capabilities of SAP54 wildtype (SAP54-wt) and the truncated proteins in yeast two-hybrid (Y2H) experiments. In accordance with previous results (MacLean et al., 2014), SAP54-wt specifically interacted with the MIKC-type MTFs SEP3, APETALA1 (AP1), and AGAMOUS-LIKE6 (AGL6) from *A. thaliana*, but not with their paralogs APETALA3 (AP3), PISTILLATA (PI) and AGAMOUS (AG) (Figures 3b and c). Deletion of either one of the two putative helices resulted in SAP54 completely losing its interaction capability (Figures 3b and c). In contrast, deletion of the predicted turn in the center of the protein did not change its interaction behavior, which would be consistent with the hypothesized modular domain architecture of SAP54 (Figures 3b and c). In addition to the deletions mentioned above, we also truncated the C-terminal 11 amino acids of the second α-helix (amino acids 81-91). Interestingly, this SAP54Δ81-91 exhibited failure to interact, suggesting the presence of an essential interaction site for MIKC-type MTFs in this region. Immunoblotting analyses (western blot) revealed that the truncated SAP54 proteins are expressed in yeast (Figure S2), indicating that the absence of yeast growth in the abovementioned Y2H studies can be attributed to the failure of the respective SAP54 derivatives to interact with MIKC-type MTFs, rather than to their absence.

**Figure 3.**
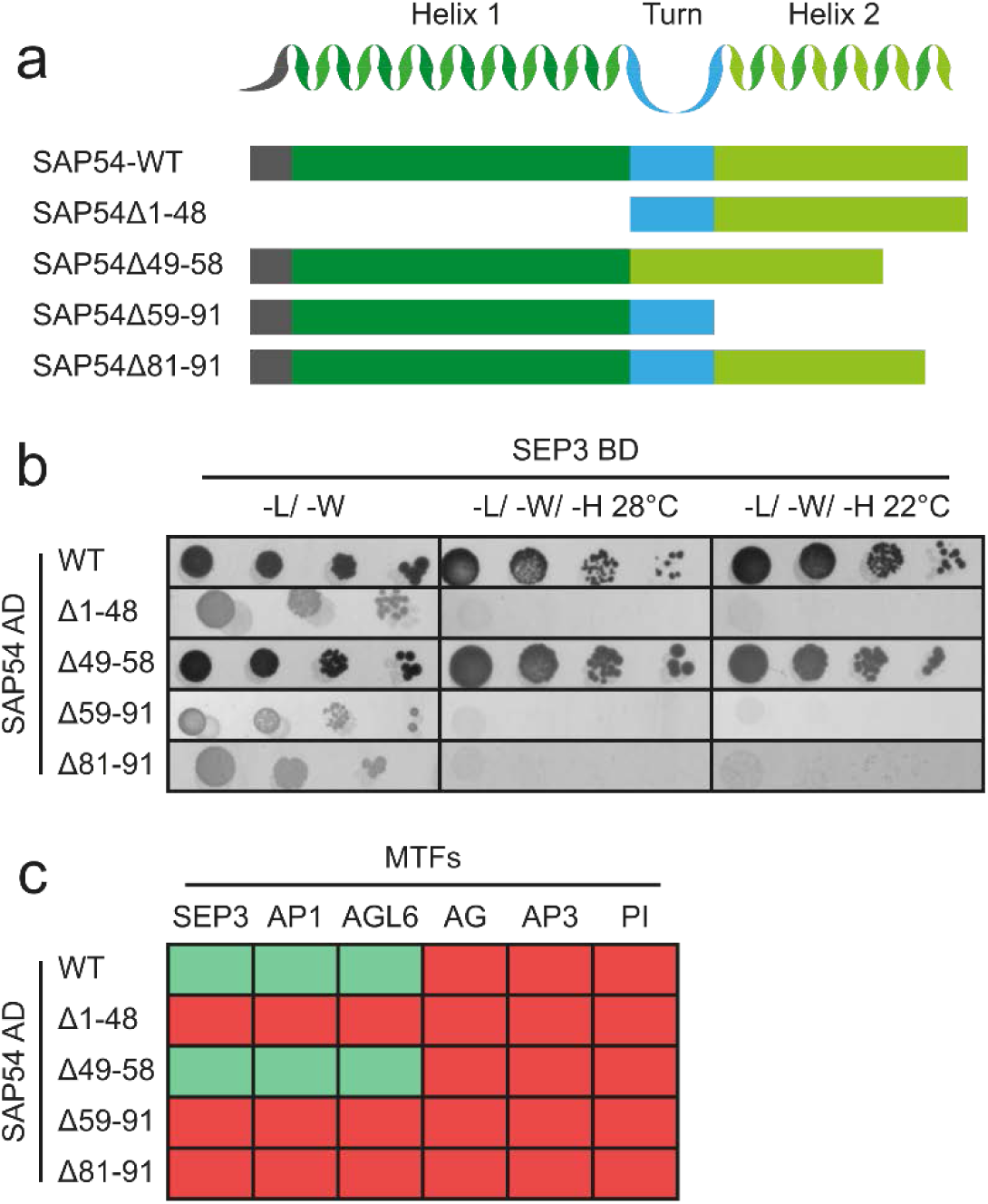
Interaction of truncated SAP54 mutants with MIKC-type MTFs in Y2H. (a) Schematic of the deletion constructs. The presumably unstructured N-terminus, the first predicted helix, the putative turn region, and the second predicted helix are depicted in gray, dark green, blue and light green, respectively. (b) Y2H interaction screen of SAP54-wt and truncation mutants fused to the activation domain (AD) against SEP3 fused to the binding domain (BD). Normal growth of yeast was controlled on SD-Leu-Trp plates (-L/-W). Interactions were tested on SD-Leu-Trp-His plates with 3 mM 3-amino1, 2, 4-triazole (-L/-W/-H) at two different temperatures (28 °C and 22 °C) and in four dilutions (1:10, 1:100, 1:1000, and 1:10000). (c) Summary of Y2H results for SAP54-wt and truncation mutants against SEP3, AP1, AGL6, AG, AP3, and PI. Each interaction was tested twice in both directions. A combination was scored as ‘interacting’ (green) if yeast growth was observed for at least one direction.

### Helix breaking point mutations impair the interaction ability of SAP54

To further analyze the importance of the α-helical elements of SAP54 for supporting interaction with MIKC-type MTFs, single amino acids at positions ‘a’ and ‘d’ of the heptad repeat of the proposed helices were substituted by proline residues. Proline was chosen because it carries helix-breaking properties due to its cyclic side chain structure, which causes steric conflicts in a helical chain conformation (Nilsson et al., 1998; Richardson, 1981). We created four individual proline substitution mutants for L27, I45, L65 and I69, respectively. Whereas L27, L65, and I69 are located close to the center of putative helix one and two, respectively, the substitution I45 is located at the C-terminus of the first putative helix and close to the predicted turn region (Figure 1a). CD spectra were recorded for purified SAP54-L27P, -I45P, and -L65P as described above to determine their respective secondary structure content. As expected, mutants SAP54-L27P and -L65P presented with a significant reduction of α-helical content (Figure 2a and Table 1). In case of the L65P substitution the formation of helical structures was almost completely abolished and analysis of SAP54-L65P in the ‘Uversky plot’ indicated that the protein was probably entirely unfolded (Figure 2b). Interestingly, the substitution of isoleucine by proline at position I45 (I45P) showed the mildest reduction of α-helical content (Table 1). Y2H analyses revealed that SAP54-L27P, -L65P, and -I69P, all expressed well in yeast, were unable to interact with any of the investigated MIKC-type MTFs (Figures 4a and b). In contrast, SAP54-I45P was still able to interact with SEP3 and AP1, whereas the interaction with AGL6 was lost.

**Figure 4.**
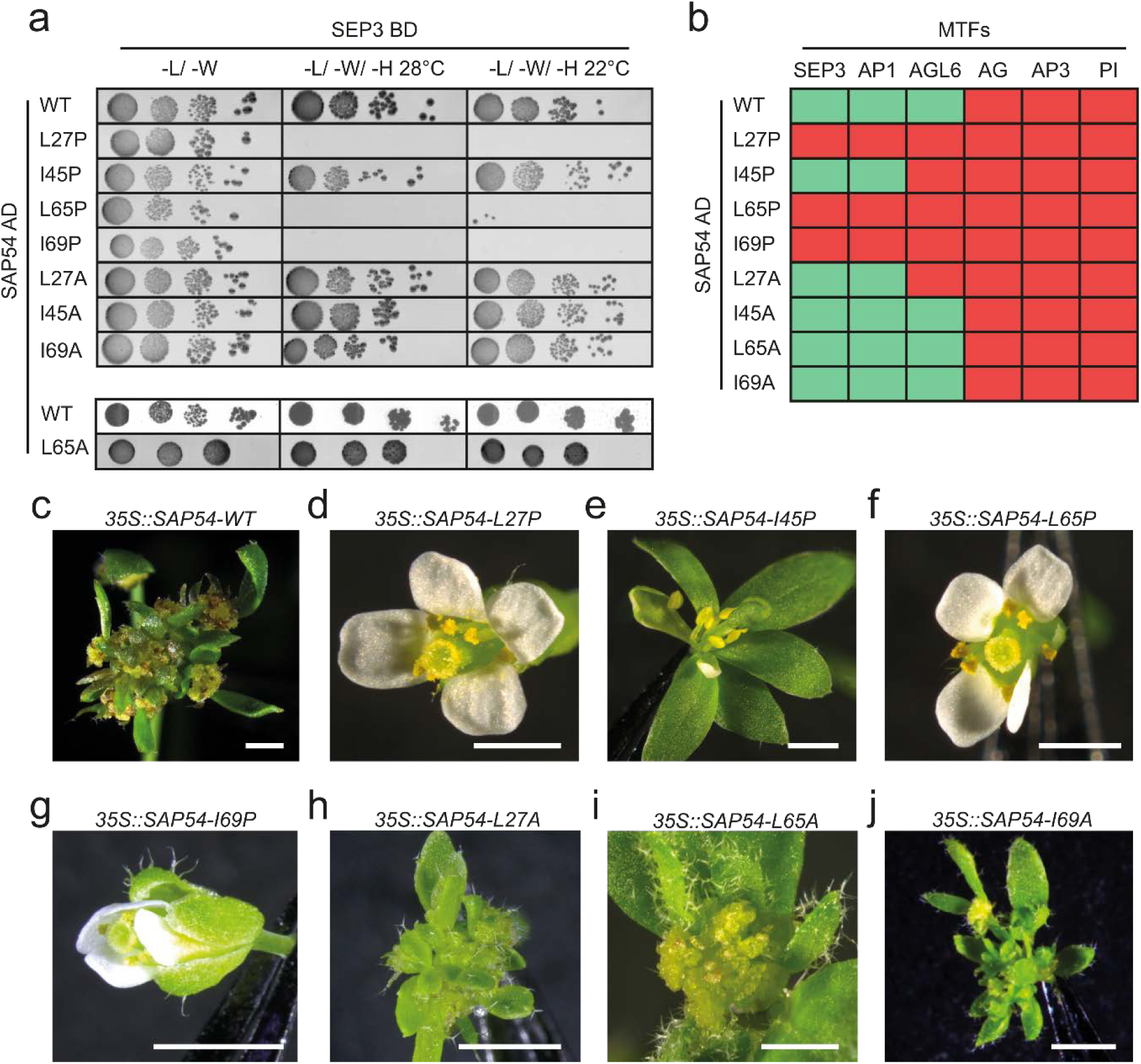
Interaction capabilities and overexpression phenotypes of SAP54-wt and single amino acid substitution mutants. (a) Y2H interaction screen of wild type and mutated versions of SAP54 fused to the activation domain (AD) against SEP3 fused to the binding domain (BD). Normal growth of yeast was controlled on SD-Leu-Trp plates (-L/-W). Interactions were tested on SD-Leu-Trp-His plates with 3 mM 3-amino1, 2, 4-triazole (-L/-W/-H) at two different temperatures (28 °C and 22 °C) and in four dilutions (1:10, 1:100, 1:1000, and 1:10000). Another SAP54-AD WT control for SAP54-AD-L65A was added due to a different time point of experimental procedure. (b) Summary of Y2H results for SAP54-wt and single amino acid substitution mutants against SEP3, AP1, AGL6, AG, AP3, and PI. Each interaction was tested twice in both directions. A combination was scored as ‘interacting’ (green) if yeast growth was observed for at least one direction. (c - j) Floral phenotypes of *A. thaliana* plants overexpressing SAP54-wt and single amino acid substitution mutants under control of the CaMV 35S-promotor. Scale bar depicts 1 mm.

In order to distinguish whether the altered interaction behavior of the proline substitution mutants was indeed related to the helix breaking properties of proline or if the substitution sites simply constitute crucial interaction sites, we additionally substituted all investigated amino acid positions by alanine (L27A, I45A, L65A and I69A). Similar to leucine and isoleucine, alanine has a high helical propensity and would thus not be expected to interfere with helix formation (Kukenshoner et al., 2014; Mason et al., 2009; Moitra et al., 1997). However, due to its small side chain, substitutions to alanine would abrogate specific residue-residue interactions at the respective sites (Cunningham and Wells, 1989; Morrison and Weiss, 2001). In contrast to the respective proline substitutions, all alanine substitution mutants were still able to interact with the known MIKC-type MTF targets of SAP54 (Figures 4a and b). Solely SAP54-L27A lost the ability to interact with AGL6 (Figure 4b). Western blot analyses revealed that all SAP54 mutant proteins had been produced in full-length in yeast (Figure S2), demonstrating that absence of yeast growth in Y2H studies was due to the inability of respective SAP54 derivatives to interact with MIKC-type MTFs, not due to their absence.

### SAP54 mutants with disturbed helix formation are unable to cause disease phenotypes

To investigate the importance of the α-helical structure of SAP54 for its ability to cause disease phenotypes, we overexpressed SAP54-wt and mutant versions under the control of the Cauliflower Mosaic Virus (CaMV) 35S promoter in *A. thaliana* Columbia-0 (Col-0). In accordance with previous results (MacLean et al., 2014) overexpression of SAP54-wt caused phyllody, i.e. the homeotic transformation of floral organs into vegetative leaf-like structures, and a partial loss of floral determinacy (Figure 4c). Furthermore, we observed clusters of undifferentiated cells arising from inflorescence meristems - a phenotype partially resembling the cauliflower-like appearance of a *cauliflower* (*cal*), *ap1* double mutant (Kempin et al., 1995). When *SAP54-L27P*, *SAP54-L65P*, and *SAP54-I69P* were overexpressed, which encode for proline substitutions close to the center of the putative helices, no phenotypic alterations were observed (Figures 4d, f, and g). In contrast, floral abnormalities were observed when overexpressing *SAP54-I45P* that encodes for a proline substitution close to the putative turn region, although the phenotypic alterations were weaker than those for SAP54-wt (Figure 4e). Plants overexpressing *SAP54-I45P* developed flowers in which most floral organs were replaced by leaf-like structures; however, some flowers developed a few petals and stamens (Figure 4e). Overexpression of the alanine substitution mutants *SAP54-L27A*, *SAP54*-*L65A*, and *SAP54*-*I69A* resulted in mutant phenotypes similar to those observed for plants overexpressing *SAP54-wt* (Figures 4h - j). To ensure that absence of phenotypic alterations was due to the inability of respective SAP54 derivatives to interact with MIKC-type MTFs, not due to their absence, expression of SAP54 proteins *in planta* was confirmed by western blot analysis (Figure S2).

### SAP54 preferentially targets multimers of MIKC-type MTFs

Knowing about the high structural similarity between SAP54 and the K-domain of MIKC-type MTFs we aimed to investigate the stoichiometry of a SAP54/K-domain complex. Therefore, we performed dynamic light scattering analyses for purified SAP54 in presence of purified K-domain of SEP3 (SEP3^75-178^) for different molar ratios. SAP54 in absence of SEP3 K-domain produces single peaks at an average hydrodynamic radius of 2.17 ± 0.12 nm, resulting in an average molecular mass estimate of ∼20 ± 2 kDa, which suggests that SAP54 forms homodimers at a concentration of 50 μM under the given phosphate buffer conditions (Figure 5a, Table 2).

**Figure 5.**
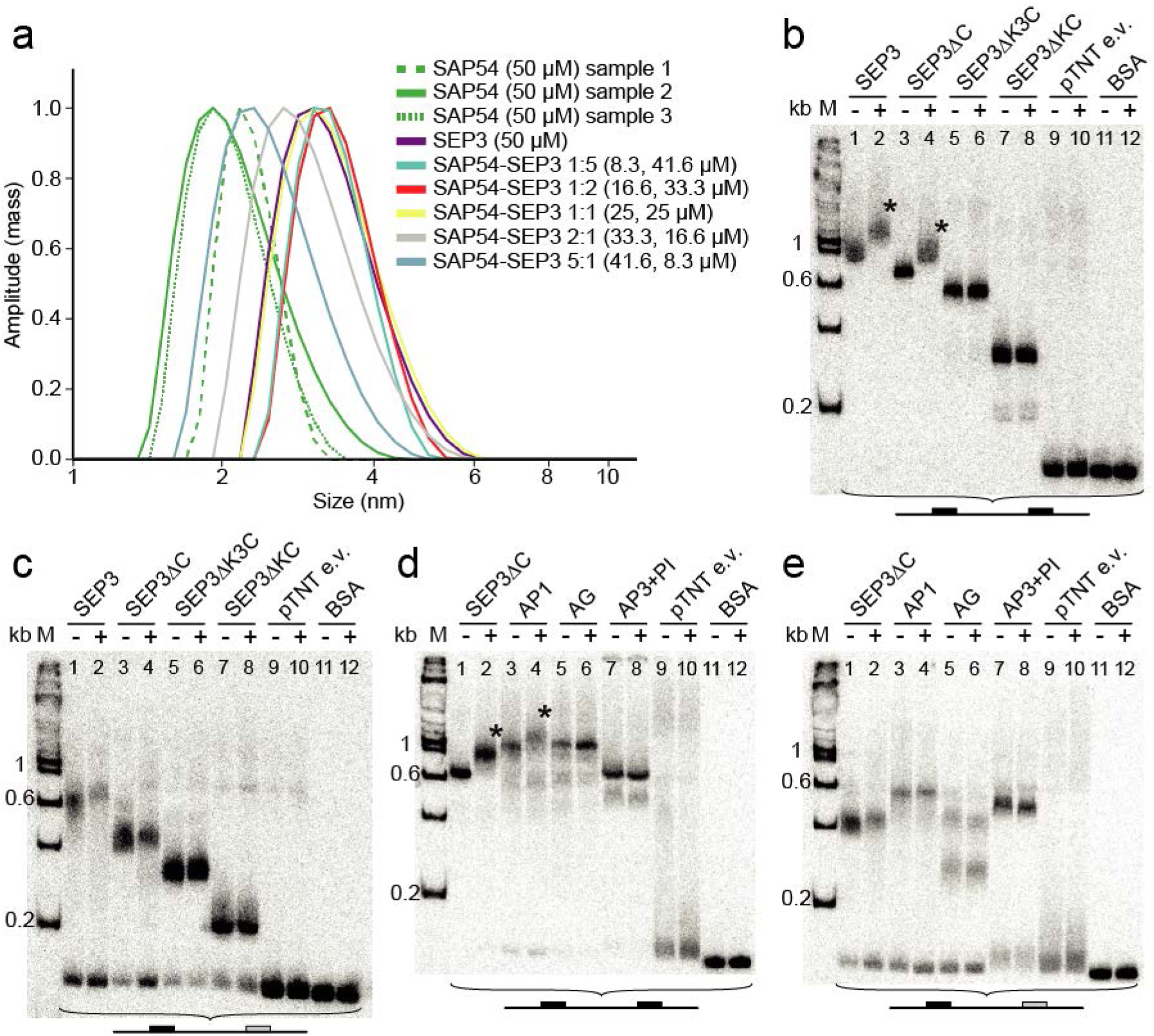
Dynamic light scattering and EMSA experiments to investigate the stoichiometry of SAP54/MIKC-type MTF complexes. (a) Dynamic light scattering plot (mass-weighted scattering intensity distribution plotted against the size of the hydrodynamic radius in nm) of SAP54, SEP3^75-178^ and mixtures of both proteins at different molar ratios. Molarities are based on the monomeric state of the proteins and were determined spectrophotometrically at 280 nm with extinction coefficients calculated using ProtParam (www.expasy.org/protparam/). For SAP54 three biological replicates were measured. (b) EMSA in which i*n vitro* translated SEP3 full length protein and truncated versions were co-incubated with a radioactively labeled DNA probe carrying two MIKC-type MTF binding sites (CArG-boxes). In lanes labeled with ‘+’ 1 μl purified SAP54 (13.3 μM) was added to the binding reaction. In lanes labeled with ‘-’ 1 μl BSA (10 mg/ml) were added instead. As negative controls, the empty pTNT vector was used as template for the *in vitro* transcription/translation (lanes ‘pTNT e.v.’) or 30 μg BSA were added instead of *in vitro* translated protein (lanes ‘BSA’). For size comparison, a radioactively labeled DNA ladder (100 bp DNA Ladder, New England BioLabs) was applied (lane M). (c) Same setup as in (b) using a DNA probe that carries only one CArG-box sequence. (d) Same setup as in (b) using the MIKC-type MTFs SEP3ΔC, AP1, AG and the obligate heterodimer AP3/PI together with a DNA probe carrying two CArG-box sequences. (e) Same setup as in (d) using a DNA probe that carries only one CArG-box sequence. Asterisks (*) depict supershifts caused by addition of SAP54.

**Table 2.**
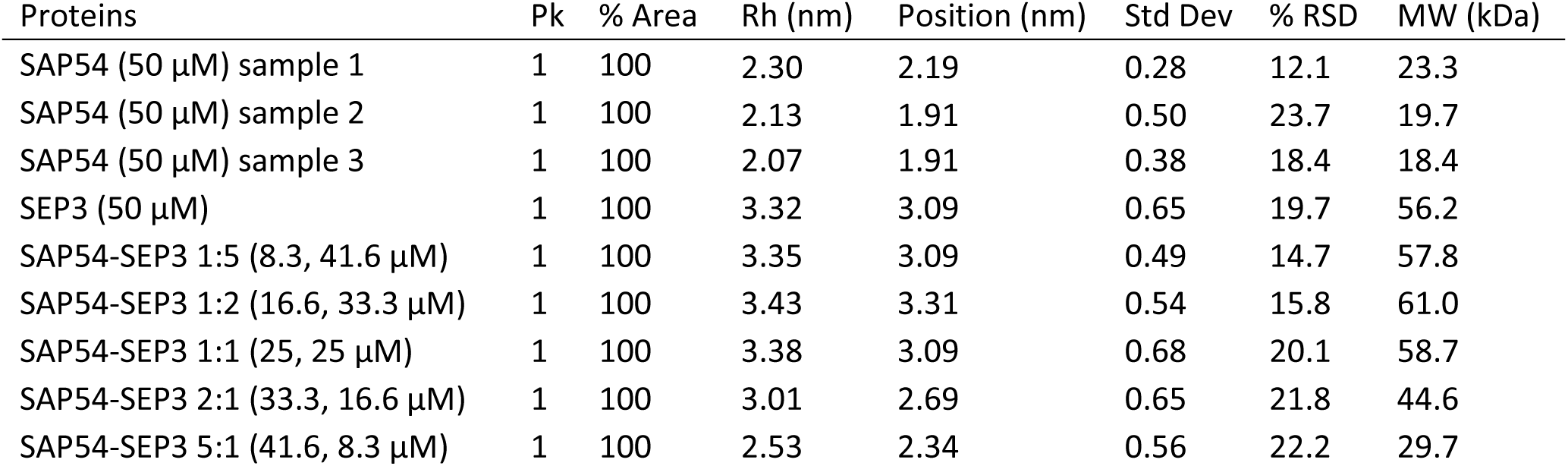
Dynamic light scattering results of SAP54, SEP3^75-178^ and mixtures of both proteins at different molar ratios. Molarities are based on the monomeric state of the proteins and were verified spectrophotometrically at 280 nm with extinction coefficients as derived from ProtParam (www.expasy.org/protparam/). Pk, number of peaks; % Area, area occupied by this major protein population; Rh (nm), hydrodynamic radius; Position, size of the hydrodynamic radius at which peak center occurs; Std, standard deviation of Rh for this peak; % RSD, percent relative standard deviation of Rh for this peak; MW (kDa), approximate molecular weight. For SAP54 three biological replicates were measured.

| Proteins | Pk | % Area | Rh (nm) | Position (nm) | Std Dev | % RSD | MW (kDa) |
| --- | --- | --- | --- | --- | --- | --- | --- |
| SAP54 (50 μM) sample 1 | 1 | 100 | 2.30 | 2.19 | 0.28 | 12.1 | 23.3 |
| SAP54 (50 μM) sample 2 | 1 | 100 | 2.13 | 1.91 | 0.50 | 23.7 | 19.7 |
| SAP54 (50 μM) sample 3 | 1 | 100 | 2.07 | 1.91 | 0.38 | 18.4 | 18.4 |
| SEP3 (50 μM) | 1 | 100 | 3.32 | 3.09 | 0.65 | 19.7 | 56.2 |
| SAP54-SEP3 1:5 (8.3, 41.6 μM) | 1 | 100 | 3.35 | 3.09 | 0.49 | 14.7 | 57.8 |
| SAP54-SEP3 1:2 (16.6, 33.3 μM) | 1 | 100 | 3.43 | 3.31 | 0.54 | 15.8 | 61.0 |
| SAP54-SEP3 1:1 (25, 25 μM) | 1 | 100 | 3.38 | 3.09 | 0.68 | 20.1 | 58.7 |
| SAP54-SEP3 2:1 (33.3, 16.6 μM) | 1 | 100 | 3.01 | 2.69 | 0.65 | 21.8 | 44.6 |
| SAP54-SEP3 5:1 (41.6, 8.3 μM) | 1 | 100 | 2.53 | 2.34 | 0.56 | 22.2 | 29.7 |

SEP3^75-178^ alone shows one peak that corresponds to a single protein population with a hydrodynamic radius of 3.32 nm resulting in an estimated molecular mass of ∼56 kDa. According to previous results (Puranik et al., 2014) the observed peak very likely represents a SEP3^75-178^ homotetramer, although the expected mass of a tetramer would be slightly lower (∼48 kDa). If SAP54 and SEP3^75-178^ were mixed in molar ratios of 1:5, 1:2, and 1:1, single peaks at 3.35, 3.43, and 3.38 nm, respectively, were observed corresponding to almost identical molecular masses of ∼58, ∼61, and ∼59 kDa (Figure 5a and Table 2). If the amount of SAP54 was increased further to molar ratios of 2:1 and 5:1, single peaks at 3.01 nm (corresponding to ∼45 kDa) and 2.53 nm (∼30 kDa) were observed (Figure 5a and Table 2). The potential influence of complex shape on the mass estimate together with the similar size of SAP54 and SEP3^75-178^ and thus the potential co-existence of complexes of very similar mass, do not allow to determine the exact stoichiometry of the observed complexes. However, according to our data it appears likely that up to a molar ratio of 1:1, SAP54 and SEP3^75-178^ form complexes that consist of 4 to 5 proteins, whereby the ratio of SAP54 and SEP3^75-178^ proteins within these complexes are probably variable. Higher concentrations of SAP54 seem to break up these complexes and instead apparently favor the formation of heterotrimers of varying composition.

To test whether SAP54 indeed tends to bind to multimers of MIKC-type MTFs, at least at low ratios of SAP54 to MIKC-MTFs, we modified a previously established electrophoretic mobility shift assay (EMSA) in order to conduct supershift experiments. If *in vitro* translated SEP3 full length protein was co-incubated with a radioactively labeled DNA probe that carries two MIKC-type MTF DNA-binding sites (so called CArG-boxes), a single retarded fraction appeared close to the 1 kb fragment of the DNA marker (Figure 5b, lane 1). As has been demonstrated in several previous studies, this fraction constitutes a SEP3 tetramer bound to both DNA-binding sites via looping the DNA in between both binding sites (Jetha et al., 2014; Melzer et al., 2009; Rümpler et al., 2018). When purified SAP54 was added to the binding reaction a supershift was observed (Figure 5b, lane 2). Since SAP54 does not bind to DNA alone (Figure 5b, lanes 10 and 12), this demonstrates that SAP54 interacts with DNA bound SEP3. Additional EMSA experiments with variable amounts of SEP3 and SAP54 indicate that SEP3 still binds cooperatively to both CArG-boxes even in a complex with SAP54 (Figure S3). Thus, it appears likely that the observed supershift fraction constitutes a DNA-bound SEP3 tetramer bound by at least one SAP54 protein. A SAP54 induced supershift could also be observed if SEP3ΔC (amino acids 1-188), which lags the C-terminal domain of SEP3, was used for the binding reaction (Figure 5b, lanes 3 and 4). In contrast, no supershift was observed if in addition to the C-terminal domain also the K3-subdomain (SEP3ΔK3C, amino acids 1-152) or the complete K-domain of SEP3 had been removed (SEP3ΔKC, amino acids 1-92), indicating that the K3-subdomain of SEP3 is necessary for the interaction with SAP54 (Figure 5b, lanes 5-8).

If SEP3 was co-incubated with a DNA probe that carries only one CArG-box a retarded fraction appeared close to the 600 bp fragment of the DNA marker (Figure 5c, lane 1). Based on previous experiments this fraction constitutes a DNA-bound SEP3 dimer (Jetha et al., 2014; Melzer et al., 2009). Interestingly, neither for DNA bound dimers of SEP3 full length protein nor for any of the truncated versions a SAP54 induced supershift was observed (Figure 5c, lanes 2-8). This strongly supports the observations made by dynamic light scattering experiments and suggests that SAP54 indeed preferentially binds to multimers of SEP3 rather than monomers or dimers. Similar results were obtained for the known SAP54 target AP1 (Figures 5d and e, lanes 3 and 4), whereas no SAP54 induced supershifts were observed for AG and AP3/PI (Figures 5d and e, lanes 5-8).

## Discussion

Employing sequence-based computer predictions and educated guesses we have previously suggested a structure for the phytoplasma effector protein SAP54. It comprises two α-helices connected by a short interhelical region in which a conserved proline breaks the α-helix (Rümpler et al., 2015). The work reported here strongly corroborates our prediction. More comprehensive and detailed *in silico* secondary structure predictions confirmed that SAP54 is composed of (at least) two putative α-helices that are separated by a turn or bend located in the centre of the protein. According to 3D structure predictions SAP54 folds into at least two coiled-coils that are separated by an interhelical region (Figure 1), very similar to the structure that had been previously hypothesized (Rümpler et al., 2015). In line with this, circular dichroism (CD) spectroscopy of purified protein revealed that SAP54 has an α-helical content of ∼71 % under our standard condition (Table 1). Only minor irregular structure features were found, most likely representing the internal turn and possibly the N-terminal end of the protein.

When either of the two putative α-helices was removed, SAP54 completely lost its ability to interact with its specific MIKC-type MTF interaction partners in Y2H experiments (Figure 3). In case of the C-terminal α-helix even deletion of only the last 11 amino acids sufficed to abolish the interaction capability completely (Figure 3). In contrast, deletion of the predicted turn in the center of the protein did not result in an altered interaction behavior (Figure 3). These findings reveal that the two putative α-helical regions are required for the interaction of SAP54 with MIKC-type MTFs, but that the internal turn is not; they also suggest that the distance between the helices does obviously not matter. The last 11 amino acids of SAP54 may comprise an essential binding motif for MIKC-type MTFs, but its exact role remains unclear.

To further test the functional relevance of the predicted structure we introduced single amino acid substitutions to proline and alanine at different heptad repeat ‘a’ and ‘d’ positions of the proposed helices of SAP54. When leucine or isoleucine residues near the center of the first (L27) or second (L65, I69) α-helix were substituted by the helix-breaking proline, α-helical content of the mutant proteins was indeed massively reduced (Figure 2). Y2H analyses revealed that the mutant proteins were unable to interact with any of the investigated MIKC-type MTFs (Figure 4). Most importantly, in contrast to the wild-type SAP54 these mutant proteins were not able anymore to cause disease phenotypes such as phyllody when expressed in *A. thaliana* (Figure 4). To demonstrate that changes in SAP54 features of the proline substitution mutants was due to the helix-breaking property of proline rather than simply the mutation of crucial interaction sites, we substituted the same residues also with alanines, which do not break helices but may interfere with the interaction of SAP54 with other proteins. However, the respective mutant SAP54 proteins were still able to interact with almost all the tested MIKC-type MTF targets of the wild-type protein in Y2H experiments, and were still capable to cause disease symptoms when expressed in *A. thaliana* (Figure 4). Interestingly, a substitution of isoleucine to proline at position near the C-terminal end of the first putative helix (I45P) showed only a moderate reduction of α-helical content (Figure 2). In line with this mild effect the interaction with only one of the investigated MIKC-type MTF was abolished (Figure 4), and, accordingly, also some ability of the mutant protein in causing phyllody was retained, even though compared to the wild-type protein its activity was clearly reduced (Figure 4).

Taken together, our findings suggest a strong causal link between the α-helical structure of major parts of SAP54, its ability to interact with some MIKC-type MTFs in Y2H analyses, and its function as an effector protein that causes homeotic transformation of floral organs *in planta*. All in all we consider our data as a strong support for our previous hypothesis on the structure of the SAP54 protein and its functional relevance (Rümpler et al., 2015).

While this manuscript was in preparation, Iwabuchi et al. (2019) published the crystal structure of PHYLLOGEN1_OY_ (PHYL1_OY_), a homolog of SAP54 from *Candidatus* Phytoplasma asteris strain Onion Yellows (OY), by X-ray diffraction. Likewise, Liao et al. (2019) published the crystal structure PHYL1_PnWB_ from *Candidatus* Phytoplasma strain Peanut Witches’ Broom. PHYL1_OY_ shares about 87 % sequence identity with SAP54 (Figure 1a) and generates similar symptoms *in planta*, suggesting that both proteins work in very similar ways (Maejima et al., 2014). In contrast, PHYL1_PnWB_ shares only about 53% sequence identity with SAP54 and is hence probably more distantly related than PHYL1_OY_ (Figure 1a; Liao et al., 2019). Nevertheless, the crystal structures of both proteins are very similar, comprising two α-helices connected by a central linker; they thus corroborate all our conclusions about SAP54 outlined here.

Moreover, Iwabuchi et al. (2019) also reported that mutational changes in the α-helices resulted in the loss of phyllody-inducing activity. In contrast to the effects of single amino acid substitutions studied by us, Iwabuchi et al. (2019) investigated the consequences of the insertion of two amino acids in a row in order to gain rotational shifts in the axes of the α-helices. While insertions in either the first or second helix abolished the interaction with some MIKC-type MTFs (SEP1 – SEP4), the degradation of SEP3, and the ability to induce disease symptoms (phyllody), insertion in the linker region had no such effects. Likewise, Liao et al. (2019) reported that some double amino acid substitutions in both either the first or second helix of PHYL1_PnWB_ abolished the interaction with the K-domain of SEP3 in Y2H experiments. These findings, summarized in Figure 6 strongly corroborate all structure-function relationships involving SAP54 outlined above.

**Figure 6.**
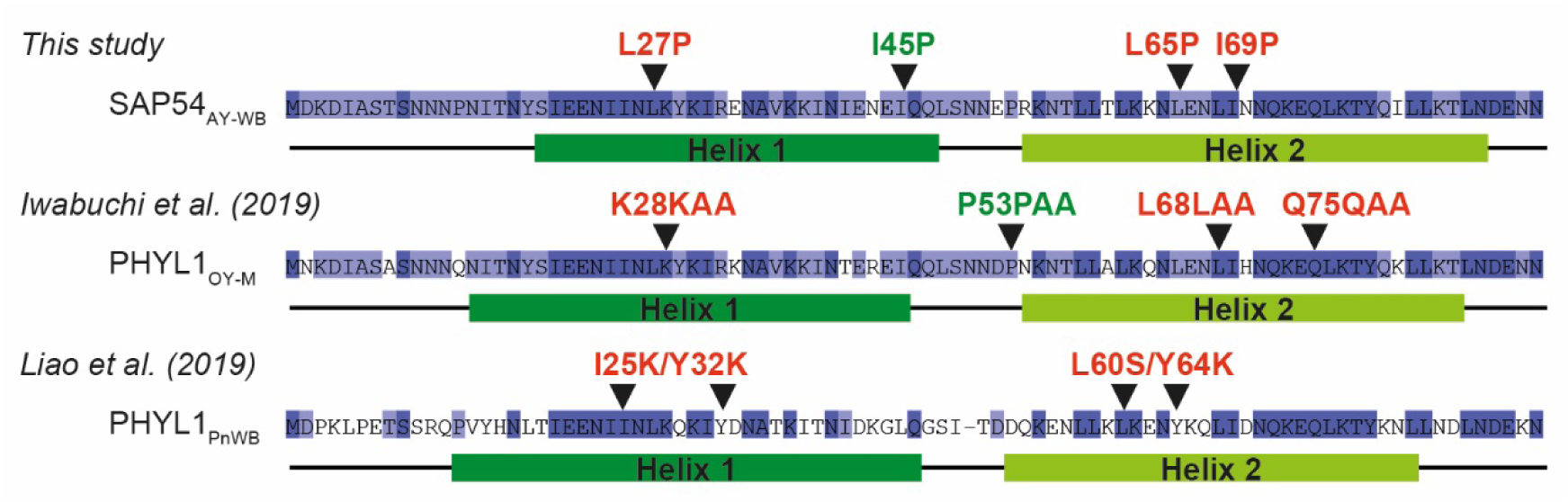
Comparison of the results of this study, Iwabuchi et al. (2019) and Liao et al. (2019). Aligned amino acid sequences of SAP54 from *Candidatus* phytoplasma asteris strain Aster Yellows Witches’ Broom, PHYL1_OY_ from *Candidatus* phytoplasma asteris strain Onion Yellows and PHYL1_PnWB_ from *Candidatus* phytoplasma Peanut witches’-broom phytoplasma. Conserved amino acid positions are marked by blue background color. Single amino acid substitutions to proline of SAP54 performed in this study, double alanine insertions of PHYL1_QY_ performed in Iwabuchi et al. (2019) and double amino acid substitutions of PHYL1_PnWB_ performed in Liao et al. (2019) are depicted by black triangles. SAP54/PHYL1_OY_ mutants that did not interact with known MIKC-type MTF targets in Y2H and that were unable to induce phyllody symptoms when overexpressed in *A. thaliana* are color coded in red, whereas mutants that were still able to induce phyllody are shown in green. Both PHYL1_PnWB_ double amino acid substitution mutants are color coded in red as they showed no interaction with the K-domain of SEP3 in a His-pulldown assay, although PHYL1_PnWB_-I25KY32K to a certain extent was still able to mediate degradation of SEP3, if both proteins were co-expressed in *Nicotiana benthamiana*.

Taken all three studies together, and considering the high sequence similarity also to other homologous proteins (Figure 1), we have now a pretty good impression about the structure of SAP54-like proteins: it comprises about 90 amino acids that fold into two α-helices connected by a small, unstructured loop. SAP54-like proteins may not have a rigid structure, however; to a certain degree it may depend on environmental conditions, and it may undergo conformational changes upon binding to and release from K-domain (Liao et al. 2019). Also the functional relevance of the structure for interaction with target proteins and for causing disease phenotypes is now well documented (this study; Iwabuchi et al., 2019). These findings will help to better understand the mode of action of SAP54-like proteins in plants.

In fact, our knowledge about the molecular mechanism of SAP54 activity is still quite limited. This applies even to the basic facts of protein-protein interactions. What, to start with, is the composite structure of SAP54 alone? Iwabuchi et al. (2019) found tetramers of SAP54 in the crystals they investigated, but since SAP54 did not interact with itself in Y2H experiments, they concluded that SAP54 exists as monomer. It is well known, however, that Y2H experiments are biased against the detection of homodimer formation (Newman et al., 2000; Smirnova et al., 1999), so this conclusion does not appear to be cogent. Liao et al. (2019) concluded from sedimentation rates during analytical ultracentrifugation that native PHYL1_PnWB_ exist as a monomer. However, in their kind of experiment they also found monomers (and tetramers) of K-domain fragments of SEP3, whereas in other experiments only dimers and tetramers had been detected (Puranik et al., 2014). The conditions during analytical ultracentrifugation thus may have promoted the dissociation of protein complexes. In line with this view, dimers seemed to be present in the crystals assessed by Liao et al. (2019). In fact, employing dynamic light scattering, we found complexes with an apparent molecular mass of ∼20 kDa, suggesting that SAP54 predominantly formed homodimers under our experimental conditions (Figure 5a, Table 2). However, since the primary sequence of PHYL1_PnWB_ is quite different from that of SAP54 and PHYL1_OY_ (Figure 1a) it currently cannot be completely ruled out that PHYL1_PnWB_ has a deviant interaction behavior, despite the high structural similarity of all three proteins.

Little is also known about heteromeric interactions of SAP54. It appears well established, at least in *A. thaliana*, that binding of SAP54 to the K-domain of a specific subset of MIKC-type MTFs is a prerequisite for the degradation of these proteins via the ubiquitin-proteasome pathway (UPP) (MacLean et al., 2014; Maejima et al., 2014). This binding likely depends on sequence and structural similarity between SAP54 and the K-domain, which both fold into α-helices and contain heptad repeats facilitating the formation of coiled-coils, as previously suggested (Rümpler et al., 2015). The similarity between SAP54 and the K-domain of MIKC-type MTFs may have originated by convergent evolution during which SAP54 achieved molecular mimicry of the K-domain (Rümpler et al., 2015). More studies on the sequence and structure of SAP54-like proteins in a phylogenetic context are required in the future to clarify the case, but are potentially hampered by frequent horizontal transfer of *SAP54* genes (Rümpler et al., 2015).

For the sake of simplicity we have previously suggested that SAP54 may form heterodimeric coiled-coils with the K-domain of MIKC-type MTFs (Rümpler et al., 2015). In line with this are data obtained by cross-linking coupled mass spectrometry suggesting that PHYL1_PnWB_ and the K-domain of SEP3 dimerize via a quadruple-stranded coiled-coil interaction (Liao et al., 2019). However, according to the dynamic light scattering and EMSA results reported here, it now seems more likely that SAP54 preferentially targets multimers of MIKC-type MTFs rather than monomers or dimers (Figures 5 and S4). This observation could probably even help to explain the target specificity of SAP54. For several known targets of SAP54 (SEP1, SEP2, SEP3, and SEP4) it has been shown that they are capable of forming DNA-bound homotetramers (Jetha et al., 2014; Melzer et al., 2009; Rümpler et al., 2018), whereas AP3 and PI that are unable to form homotetramers (Melzer and Theißen, 2009; Rümpler et al., 2018) are not bound by SAP54 (MacLean et al., 2014). Furthermore, there is evidence that SAP54 binds not only to MIKC-type MTFs, but also to RAD23C and RAD23D, i.e. a subset of the RAD23 proteins in *A. thaliana* (Iwabuchi et al., 2019; MacLean et al., 2014). RAD23 is known to transfer ubiquitylated substrates to the 26S proteasome (Farmer et al., 2010), so SAP54 may just function as an adapter that connects MIKC-type MTFs with RAD23 proteins to deliver them to the proteasome (MacLean et al., 2014). The exact place and mode of RAD23 binding to SAP54 is unknown, however several models have been proposed in which MIKC-type MTFs and RAD23 bind simultaneously, or alternatively, to SAP54 (MacLean et al., 2014). α-helical coiled coils are versatile protein domains that can contain more than two α-helices in parallel or antiparallel direction (Lupas and Gruber, 2005), and RAD23 proteins have a high content of α-helical structure (Farmer et al., 2010; Hurley et al., 2006). Thus, it is conceivable that not only SAP54 and MIKC-type MTFs may form a multimeric coiled-coil structure, but probably even RAD23 can be incorporated in such a complex. Future analyses of complexes of SAP54 with K-domains and/or RAD23 proteins by nuclear magnetic resonance (NMR) or X-ray diffraction may clarify the case.

## EXPERIMENTAL PROCEDURES

### Sequence collection, multiple sequence alignment, and structure predictions

The sequence collection of *SAP54* orthologues was compiled via nucleotide BLAST searches (Altschul et al., 1990) against the NCBI nucleotide collection and whole as well as partial genome sequences of diverse phytoplasma strains (Table S2) using the CDS of *SAP54* from AY-WB as query sequence. The nucleotide sequences were translated into the encoded amino acid sequences using EXPASY translate (Gasteiger et al., 2003) and subsequently aligned with MAFFT (Katoh and Standley, 2013) applying L-INS-I mode. Secondary structure and 3D structure predictions were performed using PCOILS (Lupas et al., 1991), PSIPRED (McGuffin et al., 2000), Jpred4 (Drozdetskiy et al., 2015), SCRATCH Protein Predictor (Cheng et al., 2005), QUARK Online (Xu and Zhang, 2012), and SWISS-MODEL (Waterhouse et al., 2018).

### Cloning of *SAP54* and *SEP3^75-178^*

The coding sequences of the secreted part of *SAP54* from Aster Yellows-Witches’ Broom phytoplasma (CP000061.1, AYWB_224) and *SEP3* (NM_102272.4) K-domain (amino acids 75-178) were codon optimized for expression in *E. coli* strain K12 with JCAT (Grote et al., 2005) (http://www.jcat.de/) and synthesized by Thermo Fisher Scientific (Table S3). Restriction sites for NdeI and BamHI were introduced at the beginning and end of the coding sequences. The synthesized sequences were amplified via PCR and cloned into the pET-15b expression vector (Novagen, www.merckmillipore.com) using NdeI and BamHI restriction sites to create expression constructs with an N-terminal 6-His-Tag followed by a thrombin cleavage site (pET-15b-*SAP54*, pET-15b-*SEP3^75-178^*). Mutated versions of *SAP54* were constructed by site-directed mutagenesis (Zheng et al., 2004).

### Expression and purification of SAP54 and SEP3^75-178^

For protein expression *E. coli* Tuner (DE3) cells (Novagen, www.merckmillipore.com) were transformed with pET-15b-*SAP54* and pET-15b-*SEP3^75-178^* using heat-shock transformation. Cells were grown in Luria-Bertani (LB) medium containing 100 µg/ml ampicillin at 37°C and 180 rpm. An overnight culture was used to inoculate pre-warmed medium (1:10) which was then grown to an optical density at 600 nm (OD_600_) of 0.5. The culture was centrifuged, resuspended and added to new medium until an OD_600_ of 0.8. Protein expression was induced by addition of 0.5 mM isopropyl-beta-D-1-thiogalactopyranoside (IPTG). Cells were harvested 3h after induction by centrifugation at 5000 x g at 4°C for 20 min. For purification, cells were resuspended in lysis buffer (50 mM Tris, pH 8.0, 300 mM NaCl, 5 mM imidazole) and one tablet “cOmplete” protease inhibitor cocktail (Roche, www.sigmaaldrich.com). The cells were lysed and homogenized by ultrasonification and French Press® disintegration. The debris was centrifuged at 15.400 x g at 4 °C for 30 min. Supernatant was applied twice to a column containing a gel bed of 2 ml nickel-nitriloacetic acid agarose (Qiagen, www.quiagen.com) equilibrated with lysis buffer. The column was then washed with one column volume of lysis buffer, washing buffer (20 mM Tris, pH 8.0, 300 mM NaCl, 20 mM imidazole) and finally with high salt buffer (50 mM Tris, pH 8.0, 500 mM NaCl). Elution of the His-tagged proteins was performed using 50 mM Tris, pH 8.0, 300 mM NaCl, and 250 mM imidazole. Fractions with addition of thrombin protease (5 Units/mg protein, Merck, www.merckmillipore.com) were pooled and dialyzed against dialysis buffer (10 mM Tris, pH 8.0, 150 mM NaCl) over night at 4 °C. Success of the thrombin cleavage was verified via SDS-PAGE gel. The protein without His-Tag was concentrated using a Vivaspin 20 Ultrafiltration Unit (3 kDa molecular weight cutoff, Sartorius, www.sartorius.de). An appropriate volume was applied to a 1 ml loop of a gel filtration column (Superdex75 10/300 GL, GE Healthcare, www.gehealthcare.com) of an Äkta Avant System (GE Healthcare, www.gehealthcare.com). Protein fractions were assessed by SDS-PAGE and fractions of interest were pooled and concentrated to 100 µM.

### CD spectra collection

The purified protein was diluted to 20 µM in 20 mM sodium phosphate buffer pH 8.0, determined spectrophotometrically at 280 nm with extinction coefficients calculated by ProtParam (ε _SAP54_ and ε _SEP3_ = 4470, https://web.expasy.org/protparam/). Five accumulated slow scan (20 nm per minute with a 1 nm slid width and a response time of 2 sec) CD spectra of freshly prepared protein solutions were recorded in the range of 190 nm to 260 nm using a JASCO J-710 CD spectropolarimeter (Jasco, www.jasco.de) at 20 °C in a 1 mm quartz cuvette. Background subtraction using a blank CD spectrum of 20 mM sodium phosphate buffer, pH 8.0, was carried out. The raw spectra were processed using the CAPITO web server application (Wiedemann et al., 2013) (www.capito.nmr.leibniz-fli.de). For melting curves, the temperature was increased stepwise by 5°C from 20°C to 90°C. Temperature increase was with a slope of 1°C/ min, the sample was kept at target temperature for 5 min before the CD spectra were recorded using the same parameters as described above.

### Dynamic light scattering

Dynamic light scattering of SAP54 and SEP3^75-178^ were carried out in a Viscotek 802 DLS device (Malvern Panalytical, www.malvernpanalytical.com) equipped with a 50 mW fiber coupled 830 nm diode laser. For each protein solution, 50 simulated experiments (5s each) were recorded at 20°C in 50 µl volume. Protein concentrations were in the range of 8-50 µM in 20 mM sodium phosphate buffer, pH 8.0. Mass weighted distribution of hydrodynamic radii and a model of globular proteins were utilized to convert hydrodynamic radii to molecular weights. Data acquisition and processing was carried out using the OmniSIZE 3.0 software (Malvern Panalytical, www.malvernpanalytical.com).

### Yeast two-hybrid screen

The yeast two-hybrid (Y2H) screen was performed as described by (Wang et al., 2010) with the following deviations: one additional -Leu/-Trp/-His plate was incubated at 28 °C for each interaction series and interaction was scored after 5-7 days of incubation. Additionally, the β-galactosidase assay was performed on -Leu/-Trp plates. An interaction was scored when growth on both interaction plates (at 22 °C and 28 °C) and blue coloration of the colonies in the β-galactosidase assay was observed. For in-detail procedures consult experimental supplementary.

### Yeast protein extraction and western blot

The expression of SAP54-wt and mutant versions as well as MIKC-type MTFs in yeast cells was confirmed by protein extraction according to the protocol of Kushnirov (Kushnirov, 2000) followed by SDS-PAGE, western blot and immuno-detection using monoclonal antibodies raised in mouse against the GAL4 activation or binding domain (1:10000 dilution, Clontech, www.clontech.com) following the Matchmaker Monoclonal Antibodies User Manuel from Clontech. An antibody against α-tubulin (1:10000 dilution, abcam, www.abcam.com) was used as a loading control for protein extracts. Proteins were detected using the Roti-Lumin Kit (Carl-Roth, www.carlroth.com) and visualized in the ImageQuant LAS 4000mini (GE Healthcare, www.gehealthcare.com).

### Plant transformation and phenotyping

For *SAP54* overexpression studies in *A. thaliana* Col-0 plants, the CDS of the secreted part of *SAP54*-wt and mutant versions were cloned into the pGreen0229 vector under control of the CaMV 35S promoter. The vectors were used to transform *Agrobacterium tumefaciens* GV3101 cells (carrying the helper plasmid pSOUP) via electroporation at 1800 V using the Electroporator 2510 (Eppendorf). Agrobacteria were screened via colony PCR using primers flanking the 35S promoter region and the terminator region of the T-DNA. Correct clones were used for “floral-dipping” following the protocol of Clough and Bent (Clough and Bent, 1998). Plants were covered with plastic bags to retain moisture for 24 h. After maturity seeds were collected and sown on soil. BASTA (0,001%, Bayer, www.bayer.com) was applied via spraying to select transformed T1 seedlings. T1 plants were grown under long day conditions at 22 °C and phenotyped using the z-stack function of a Leica M205 FA microscope (Leica, www.leica.de).

### Plant Protein extraction and western blot

For protein extraction from transformed plants, 80 mg of leaf and inflorescence material were ground to a fine powder over 40 sec at 30 Hertz at 4°C using the Retsch MM 400 Mill (Thermo Fischer Scientific, www.thermofisher.com) and were then resuspended in 200 µl homogenization buffer (0.1 M Tris-HCL, pH 7.5, 10% glycerol, 5 mM EDTA, 2 mM EGTA, 2 mM PMSF, 40 mM β-mercaptoethanol, 2 µg/ml pepstatin, 2 µg/ml leupeptin). The solution was centrifuged for 3 min at 16,000 x g at 4°C and supernatant was used for western blot as described above. Primary antibody against SAP54 (raised in rabbit, kindly provided by Saskia Hogenhout, John-Innes Centre, Norwich) was used in a 1:2000 dilution.

### *In vitro* transcription/translation and electrophoretic mobility shift assay

*In vitro* transcription/translation and electrophoretic mobility shift assays (EMSA) were performed essentially as described by Melzer et al. (2009). The plasmids for *in vitro* transcription/translation of *A. thaliana* SEP3, SEP3ΔC (amino acids 1-188), SEP3ΔK3C (amino acids 1-152), SEP3ΔKC (amino acids 1-92), AP1, AP3, PI, and AG, namely pTNT-SEP3, pTNT-SEP3ΔC, pTNT-SEP3ΔK3C, pTNT-SEP3ΔKC, pSPUTK-AP1, pSPUTK-AP3, pSPUTK-PI, and pSPUTK-AG had been generated and described previously (Melzer et al., 2009; Riechmann et al., 1996). DNA probe preparation and labelling had been described previously (Melzer et al., 2009). The CArG-box sequence (5’-CCAAATAAGG-3’) used for probe preparation was derived from the regulatory intron of *A. thaliana AG*. To test for binding of SAP54 to DNA-bound MIKC-type MTF tetramers a 151-bp DNA probe was used that contained two CArG-box sequences in a distance of 63 bp, i.e. six helical turns (sequence: 5’-TCGAG GTCGG AAATT TAATT ATATT *CCAAA TAAGG* AAAGT ATGGA ACGTT CGACG GTATC GATAA GCTTG ATGAA ATTTA ATTAT ATT*CC AAATA AGG*AA AGTAT GGAAC GTTAT CGAAT TCCTG CAGCC CGGGG GATCC ACTAG TTCTA G-3’; CArG-box sequences are in italics). To test for binding of SAP54 to DNA-bound MIKC-type MTF dimers a DNA probe was used in which one of the two CArG-boxes and its flanking regions were exchanged by a random sequence of the same base composition (sequence: 5’-TCGAG GTCGA TAAAA CGGCA AGGAG AATTA TATTT TTATG ATGAA CATAT CGACG GTATC GATAA GCTTG ATGAA ATTTA ATTAT ATT*CC AAATA AGG*AA AGTAT GGAAC GTTAT CGAAT TCCTG CAGCC CGGGG GATCC ACTAG TTCTA G-3’). The composition and final concentrations of the protein-DNA binding reaction buffer were 1.6 mM EDTA, 10.3 mM HEPES, 1 mM DTT, 1.3 mM Spermidine hydrochloride, 33.3 ng/μl Poly dI/dC, 2.5 % CHAPS, 4.3 % glycerol, and 3.72 μg/μl BSA. For each binding reaction 0.1 ng labelled DNA probe were co-incubated with 3 μl of *in vitro* translated MIKC-type protein and either 1 μl purified SAP54 (13.3 μM) or 1 μl BSA (10 μg/μl) over night at 4 °C. The co-operative DNA-binding assay shown in Figure S3 was performed essentially as described by Melzer et al. (2009). A constant amount of 0.1 ng labelled DNA probe containing two CArG-boxes was co-incubated with variable amounts of *in vitro* translated SEP3ΔC, ranging from 0.05 - 2 μl. After measuring the relative signal intensities of all fractions for each lane using Multi Gauge 3.1 (Fujifilm), the ability of SEP3ΔC to co-operatively bind to two DNA-binding sites was quantified using equations as previously described (Melzer et al., 2009; Senear and Brenowitz, 1991). To examine whether SEP3ΔC still binds co-operatively to DNA in the presence of SAP54, the same assay was performed while adding 1 μl purified SAP54 (13.3 μM) to each binding reaction.

## ACKNOWLEDGEMENTS

We are indebted to Saskia Hogenhout (John-Innes Centre, Norwich) for providing SAP54 antibody. We are also grateful to Sabine Häfner (Fritz-Lipmann-Institute, Jena) for her skilful assistance in the purification of proteins. This work was supported by a fellowship from the International Leibniz Research School for Microbial and Biomolecular Interactions (ILRS Jena), which is part of the Jena School for Microbial Communication (JSMC) (to M.B.A.).

## SUPPORTING INFORMATION

**Figure S1.** Uversky plot of heated and recooled SAP54 protein.

**Figure S2.** Western blot to confirm expression of SAP54-wt and mutated proteins in yeast and in *A. thaliana*.

**Figure S3.** Test for co-operative DNA binding of SEP3ΔC and SEP3ΔC in a complex together with SAP54.

**Table S1.** Content of secondary structure features of SAP54-wt when exposed to rising temperature levels.

**Table S2:** List of phytoplasma whole and partial genome sequences that were searched for SAP54 orthologues.

**Table S3:** Codon optimized sequences of SAP54 from AY-WB and the K-domain of SEP3 from *A. thaliana*.

