## Supplementary for "Structural requirements of the phytoplasma effector protein SAP54 for causing homeotic transformation of floral organs"

**SUPPLEMENTARY DATA**

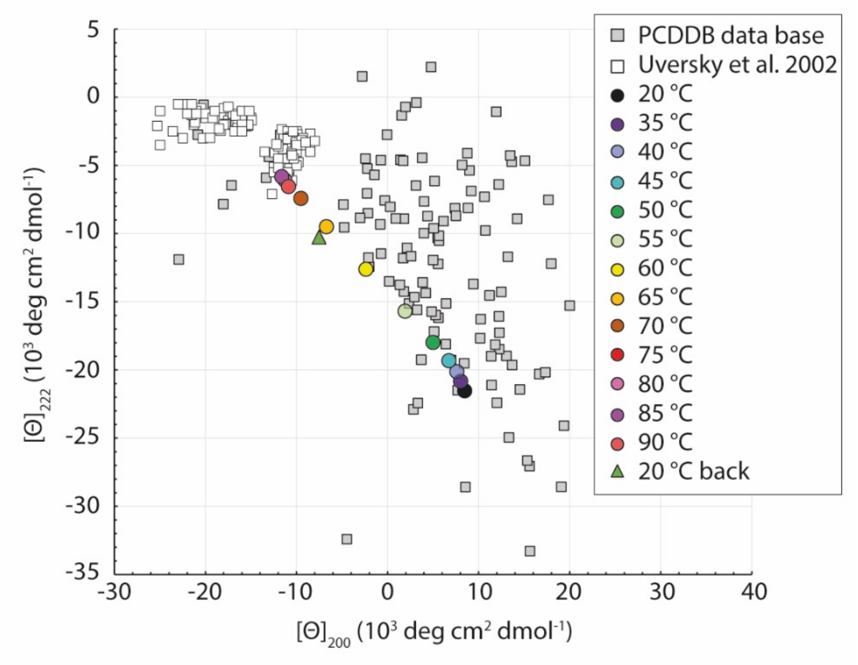


**Figure S1.** Uversky plot of heated and recooled SAP54 protein. SAP54 shows a stepwise denaturation with a melting point at approximately 56 °C. Even at 90°C, the CD spectra indicate a thermostable residual structure, rather than complete denaturation (see also Table S1). Reverting the temperature from 90°C back to 20°C (sample 20°C back) the CD analysis indicates, that the protein does not regain its native fold following heat denaturation.

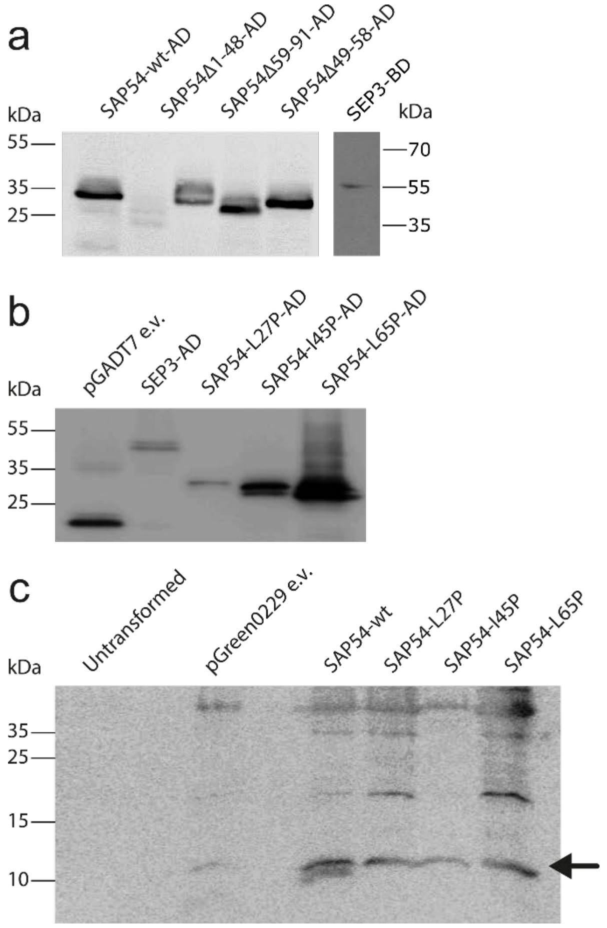


**Figure S2.** Western blot to confirm expression of SAP54-wt and mutated proteins in yeast and in *A. thaliana*. (a) Western blot of protein extracted from cultured yeast cells expressing SAP54-wt-AD and the truncated versions SAP54Δ1-48-AD, SAP54Δ59-91-AD, and SAP54Δ49-58-AD using antibodies against the GAL4 activation domain (left) and from yeast cells expressing SEP3-BD using antibodies against the GAL4 DNA-binding domain (right). (b) Western blot of protein extracted from cultured yeast cells expressing SEP3-AD and the SAP54 proline substitution mutants SAP54-L27P-AD, SAP54-I45P-AD, and SAP54-L65P-AD using antibodies against the GAL4 activation domain. The empty vector pGADT7 expressing only the activation domain was used as positive control. (c) Western blot of protein extracted from untransformed *A. thaliana* Col-0, *A. thaliana* transformed with the empty vector pGreen0229, and *A. thaliana* plants overexpressing SAP54-wt and the proline substitution mutants SAP54-L27P, SAP54-I45P, and SAP54-L65P using an antibody against SAP54. Black arrow depicts the band at the expected size of ~11 kDa.

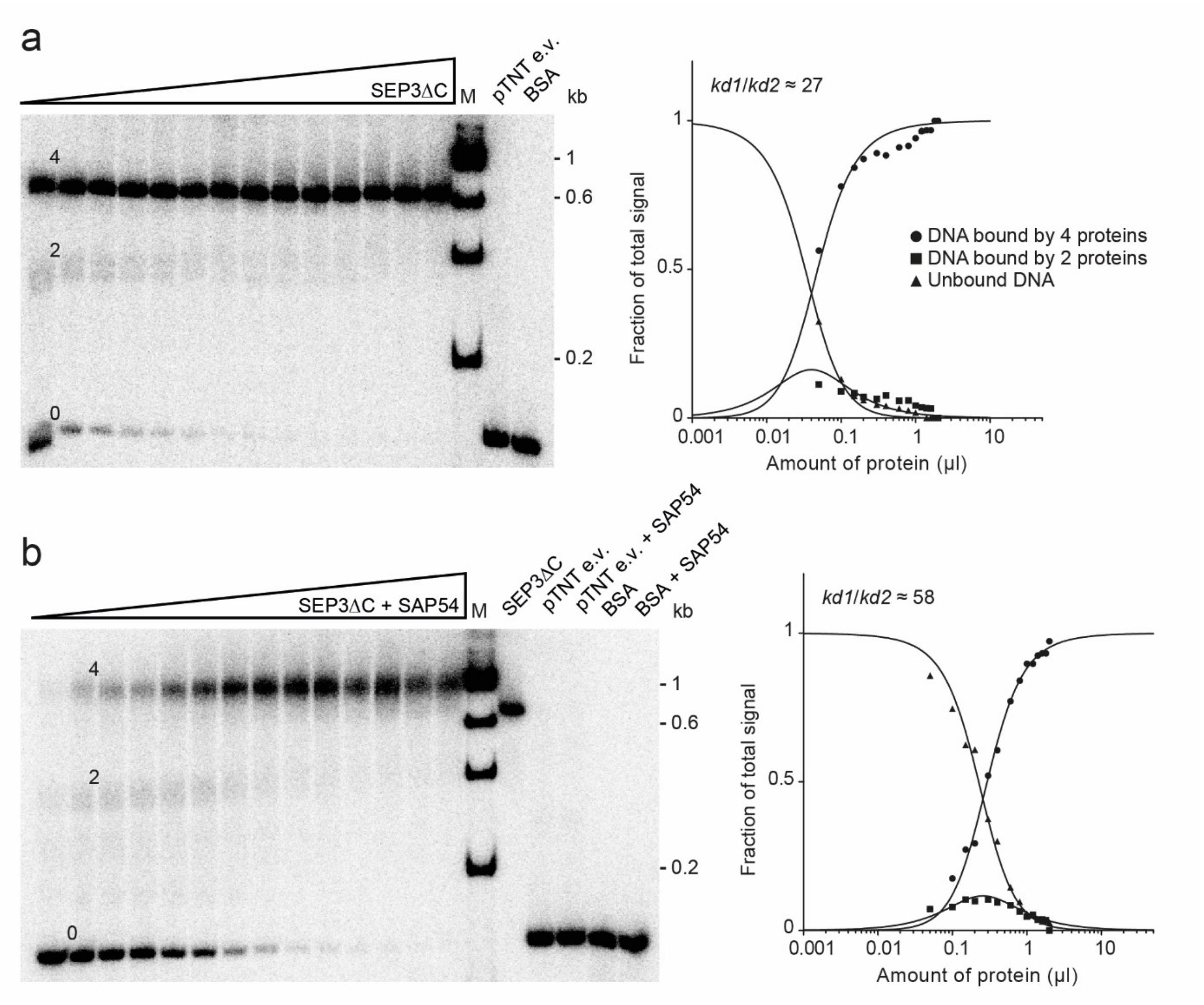


**Figure S3.** Test for co-operative DNA binding of SEP3ΔC and SEP3ΔC in a complex together with SAP54. Increasing amounts of either (a) *in vitro* translated SEP3ΔC alone or (b) *in vitro* translated SEP3ΔC together with purified SAP54 were co-incubated with 0.1 ng of a radioactively labeled DNA-probe that carried two CArG-boxes. With increasing amounts of protein three fractions of different electrophoretic mobility occur: a fraction of high electrophoretic mobility that constitutes unbound DNA (labeled with '0'), a fraction of intermediate electrophoretic mobility that constitutes a DNA probe bound by 2 SEP3ΔC proteins ('2') and a fraction of low electrophoretic mobility that constitutes a DNA probe bound by 4 SEP3ΔC proteins ('4'). As negative controls, the empty pTNT vector was used as template for the *in vitro* transcription/translation (lanes 'pTNT e.v.') or 30 µg BSA were added instead of *in vitro* translated protein (lanes 'BSA'). For size comparison, a radioactively labeled DNA ladder (100 bp DNA Ladder, New England BioLabs) was applied (lane M). To quantify the ability for co-operative DNA binding and thus the ability of SEP3ΔC to form DNA-bound tetramers the relative signal intensities of all fractions were measured for each lane. The resulting ratios were used to estimate the dissociation constant for binding of the first SEP3ΔC dimer (*kd1*) and for binding of a second SEP3ΔC dimer to a DNA fragment where one of the two CArG-boxes are already occupied (*kd2*) using equations as previously described (Melzer et al., 2009; Senear and Brenowitz, 1991). The resulting *kd1*/*kd2* values are depicted in the graphs. Similar to SEP3ΔC alone also SEP3ΔC together with SAP54 showed co-operative DNA binding.

**Table S1.** Content of secondary structure features of SAP54-wt when exposed to rising temperature levels. Starting at 30°C, temperature was raised by 5°C up to 90°C. At each level, the protein was exposed to the new temperature for 5 minutes before the spectrum was recorded. An additional spectrum was recorded after the solution was brought back to 20°C (SAP54 20°C back). The results depicted were calculated with the help of CAPITO (Wiedemann et al., 2013).

| Sample | % α-helix | % β-sheet | % irregular |
|---|---|---|---|
| SAP54 30°C | 62 | 4 | 39 |
| SAP54 35°C | 61 | 3 | 34 |
| SAP54 40°C | 58 | 1 | 35 |
| SAP54 45°C | 50 | 6 | 41 |
| SAP54 50°C | 42 | 2 | 39 |
| SAP54 55°C | 29 | 9 | 47 |
| SAP54 60°C | 23 | 10 | 52 |
| SAP54 65°C | 20 | 19 | 57 |
| SAP54 70°C | 12 | 19 | 61 |
| SAP54 75°C | 6 | 25 | 63 |
| SAP54 80°C | 17 | 18 | 64 |
| SAP54 85°C | 16 | 26 | 67 |
| SAP54 90°C | 4 | 23 | 68 |
| SAP54 20°C back | 12 | 6 | 57 |

**Table S2:** List of phytoplasma whole and partial genome sequences that were searched for SAP54 orthologues.

| 16Sr group | Strain | GenBank identifier |
|---|---|---|
| 16SrI | Aster yellows witches'-broom phytoplasma | CP000061 |
| 16SrI | 'Brassica napus' phytoplasma isolate TW1 | QGKT01000000 |
| 16SrI | 'Chrysanthemum coronarium' phytoplasma strain OY-V | BBIY00000000 |
| 16SrI | Chrysanthemum yellows phytoplasma strain CYP | JSWH00000000 |
| 16SrI | Maize bushy stunt phytoplasma strain M3 | CP015149 |
| 16SrI | Milkweed yellows phytoplasma strain MW1 | AKIL00000000 |
| 16SrI | New Jersey aster yellows phytoplasma strain NJAY | MAPF00000000 |
| 16SrI | Onion yellows phytoplasma strain OY-M | AP006628 |
| 16SrI | Wheat blue dwarf phytoplasma | AVAO00000000 |
| 16SrII | Candidatus Phytoplasma aurantifolia strain WBDL | MWKN01000000 |
| 16SrII | 'Echinacea purpurea' witches'-broom phytoplasma strain NCHU2014 | LKAC01000000 |
| 16SrII | Peanut witches'-broom phytoplasma NTU2011 | AMWZ00000000 |
| 16SrIII | Candidatus Phytoplasma pruni strain CX | LHCF00000000 |
| 16SrIII | Italian clover phyllody phytoplasma strain MA1 | AKIM00000000 |
| 16SrIII | X-disease group Phytoplasma sp. Vc33 | LLKK00000000 |
| 16SrIII | Poinsettia branch-inducing phytoplasma strain JR1 | AKIK00000000 |
| 16SrIII | Vaccinium witches'-broom phytoplasma strain VAC | AKIN00000000 |
| 16SrV | Candidatus Phytoplasma ziziphi isolate Jwb-nky | CP025121 |
| 16SrIX | Candidatus Phytoplasma phoenicium strain SA213 | JPSQ00000000 |
| 16SrIX | Candidatus Phytoplasma phoenicium strain ChiP | PUUG00000000 |
| 16SrX | Candidatus Phytoplasma mali strain AT | CU469464 |
| 16SrXI | Candidatus Phytoplasma oryzae strain NGS-S10 | JHUK00000000 |
| 16SrXI | Candidatus Phytoplasma oryzae isolate Mbita1 | LTBM01000000 |
| 16SrXII | Candidatus Phytoplasma australiense | AM422018 |
| 16SrXII | Candidatus Phytoplasma solani strain 284/09 | FO393427 |
| 16SrXII | Candidatus Phytoplasma solani strain 231/09 | FO393428 |
| 16SrXII | Candidatus Phytoplasma solani strain SA-1 | MPBG00000000 |
| 16SrXII | Strawberry lethal yellows phytoplasma strain NZSb11 | CP002548 |
| unclassified | Rice orange leaf phytoplasma strain LD1 | MIEP00000000 |

**Table S3:** Codon optimized sequences of SAP54 from AY-WB and the K-domain of SEP3 from *A. thaliana*. Codon optimization was performed with JCAT.

```
>SAP54
ATGGACAAAGACATCGCTTCTACCTCTAACAACAACCCGAACATCACCAA     50
CTACTCTATCGAAGAAAACATCATCAACCTGAAATACAAAATCCGTGAAA     100
ACGCTGTTAAAAAAATCAACATCGAAAACGAAATCCAGCAGCTGTCTAAC     150
AACGAACCGCGTAAAAACACCCTGCTGACCCTGAAAAAAAACCTGGAAAA     200
CCTGATCAACAACCAGAAAGAACAGCTGAAAACCTACCAGATCCTGCTGA     250
AAACCCTGAACGACGAAAACAACTAA

>SEP3
ATGTACGGTGCTCCGGAACCGAACGTTCCGTCTCGTGAAGCTCTGGCTGT     50
TGAACTGTCTTCTCAGCAGGAATACCTGAAACTGAAAGAACGTTACGACG     100
CTCTGCAGCGTACCCAGCGTAACCTGCTGGGTGAAGACCTGGGTCCGCTG     150
TCTACCAAAGAACTGGAATCTCTGGAACGTCAGCTGGACTCTTCTCTGAA     200
ACAGATACGTGCTCTGCGTACCCAGTTCATGCTGGACCAGCTGAACGACC     250
TGCAGTCTAAAGAACGTATGCTGACCGAAACCAACAAAACCCTGCGTCTG     300
CGTCTGGCTGACGGTTAA
```
